# A new *in vitro* assay measuring direct interaction of nonsense suppressors with the eukaryotic protein synthesis machinery

**DOI:** 10.1101/330506

**Authors:** Martin Y. Ng, Haibo Zhang, Amy Weil, Vijay Singh, Ryan Jamiolkowski, Alireza Baradaran-Heravi, Michel Roberge, Allan Jacobson, Westley Friesen, Ellen Welch, Yale E. Goldman, Barry S. Cooperman

**Affiliations:** Department of Chemistry, University of Pennsylvania, Philadelphia, PA 19104; Department of Physiology, Perelman School of Medicine, University of Pennsylvania, Philadelphia, PA 19104; Department of Biochemistry and Molecular Biology, University of British Columbia, Vancouver, BC, Canada V6T 1Z3; Department of Microbiology and Physiological Systems, University of Massachusetts Medical School, Worcester, MA 01655; PTC Therapeutics, 100 Corporate Court, South Plainfield, NJ 07080

**Keywords:** Premature termination codon (PTC), Readthrough, Ribosome, in vitro assay, eukaryotic translation, non-sense suppressors

## Abstract

Nonsense suppressors (NonSups) induce “readthrough”, i.e., the selection of near cognate tRNAs at premature termination codons and insertion of the corresponding amino acid into nascent polypeptide. Prior readthrough measurements utilized contexts in which NonSups can promote readthrough directly, by binding to one or more of the components of the protein synthesis machinery, or indirectly, by several other mechanisms. Here we utilize a new, highly-purified *in vitro* assay to measure exclusively direct nonsense suppressor-induced readthrough. Of 16 NonSups tested, 12 display direct readthrough, with results suggesting that such NonSups act by at least two different mechanisms. In preliminary work we demonstrate the potential of single molecule fluorescence energy transfer measurements to elucidate mechanisms of NonSup-induced direct readthrough, which will aid efforts to identify NonSups having improved clinical efficacy.

**Table of Contents artwork:** 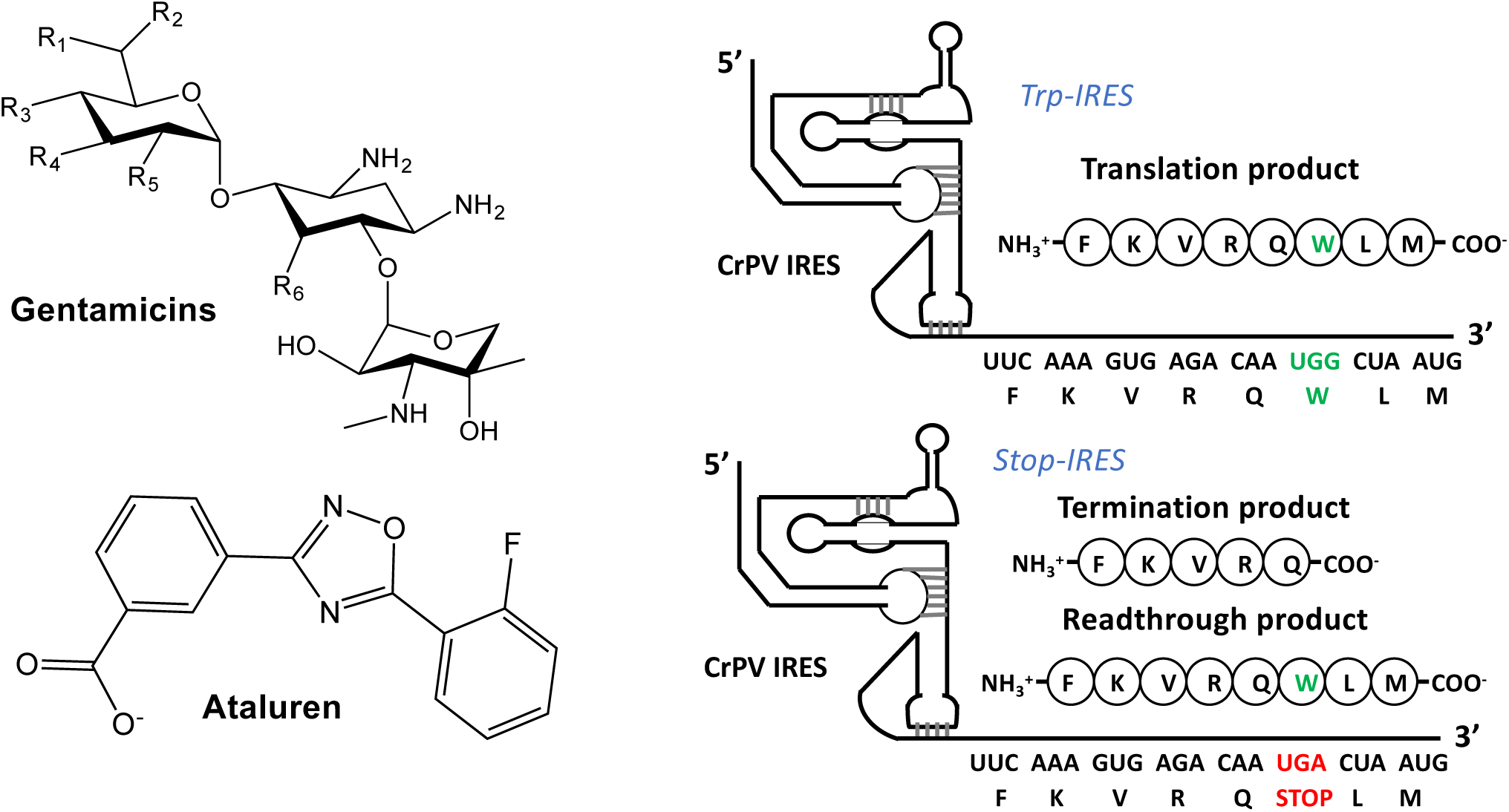

---

Premature termination codons (PTCs) arise as a consequence of nonsense mutations. Such mutations lead to the replacement of an amino acid codon in mRNA by one of three stop codons, UAA, UGA or UAG^1-3^, and result in inactive truncated protein products. Nonsense mutations constitute ~20% of transmitted or *de novo* germline mutations.^4-6^ Globally, there are ~7000 genetically transmitted disorders in humans, and ~11% of all human disease mutations are nonsense mutations.^7^ Clearly, millions of people worldwide would benefit from effective therapies directed toward PTC suppression. Clinical trials have begun to evaluate the treatment of PTC disorders with therapeutic agents called nonsense suppressors (NonSups).^8-10^ NonSups induce the selection of near cognate tRNAs at the PTC position, and insertion of the corresponding amino acids into the nascent polypeptide, a process referred to as “readthrough,” which restores the production of full length functional proteins, albeit at levels considerably reduced from wild-type. Even low rates of readthrough can improve clinical outcomes when essential proteins are completely absent. Examples of such essential proteins include cystic fibrosis trans-membrane regulator, dystrophin, and the cancer tumor suppressor proteins adenomatous polyposis coli and p53 (references provided in *Supporting Information*, Item 10).

*In vitro, ex vivo* and *in vivo* experiments and clinical trials have identified a diverse structural set of NonSups as candidates for PTC suppression therapy (Figure 1), including aminoglycosides, ataluren and ataluren-like molecules and others (references provided in *Supporting Information*, Item 10). To date, only one NonSup, ataluren (known commercially as Translarna), has been approved in the EU for clinical use, but this approval is limited to treatment of patients with nonsense-mediated Duchenne muscular dystrophy. The clinical utility of other NonSups, such as aminoglycosides, is restricted, in part, by their toxic side effects. A critical barrier to development of NonSups with broader therapeutic windows is the paucity of information regarding the precise mechanisms by which these molecules stimulate readthrough. All prior results measuring NonSup-induced readthrough of eukaryotic PTCs have been carried out using animals, intact cells or crude cell extracts. In such systems, NonSups can promote readthrough directly, by binding to one or more of the components of the protein synthesis machinery, or indirectly, either by inhibiting nonsense-mediated mRNA decay,^11^ or by modulating processes altering the cellular activity levels of protein synthesis machinery components.^12,13^ This multiplicity of possible mechanisms of nonsense suppression has complicated attempts to determine the precise mechanisms of action of specific NonSups and limited the use of rational design in identifying new, more clinically useful NonSups.

**Figure 1.**
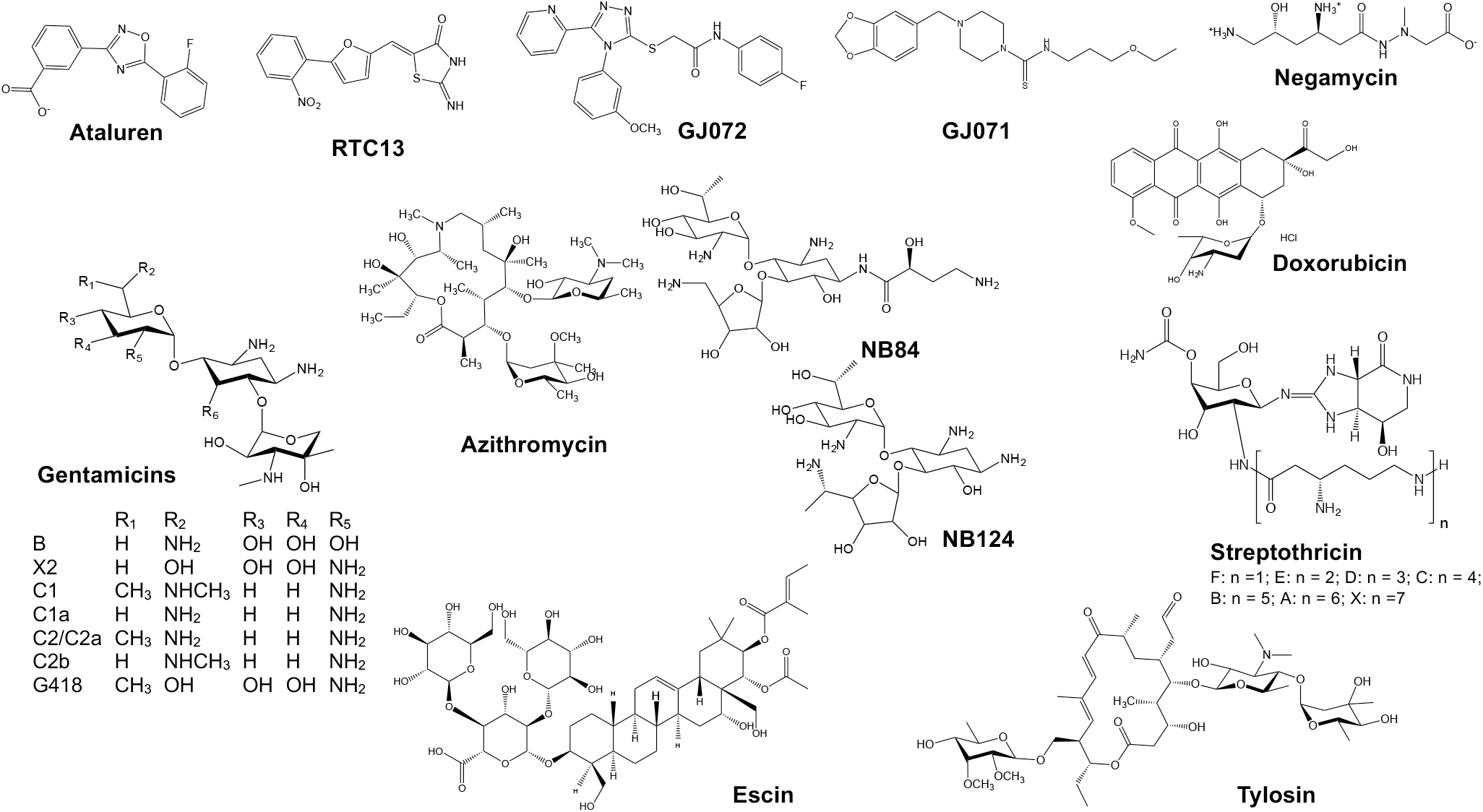
The structures of the nonsense suppressors (NonSups) studied in this work.

Recently, we developed a highly purified, eukaryotic cell-free protein synthesis system,^14^ that we apply here to distinguish NonSups acting directly on the protein synthesis machinery from those that act indirectly. We present evidence suggesting that NonSups acting directly can be divided into at least two distinctive structural groups which induce readthrough by different mechanisms. We also demonstrate the potential of using single molecule fluorescence resonance energy transfer (smFRET) to elucidate the details of such mechanisms.

Our system exploits the ability of the intergenic internal ribosome entry site (IRES) of Cricket Paralysis Virus (CrPV-IRES) to form a complex with 80S ribosomes capable of initiating the synthesis of complete proteins in cell-free assays completely lacking initiation factors.^15,16^ Structural studies (references provided in *Supporting Information*, Item 10) have shown that, prior to polypeptide chain elongation, CrPV-IRES and the closely related IRES from Taura syndrome virus occupy all three tRNA binding sites (E, P, and A) on the 80S ribosome. We recently demonstrated that the first two cycles of peptide elongation proceed very slowly due to very low rates of pseudo-translocation and translocation, but that, following translocation of tripeptidyl-tRNA, subsequent elongation cycles proceed more rapidly, presumably as a consequence of IRES removal from the ribosome.^14^

Based on these results we have now constructed an assay to directly monitor readthrough at the stop codon in the sixth position, when the faster elongation rate is well established.

For this purpose, we prepared two CrPv IRES coding sequences, Stop-IRES and Trp-IRES (Figure 2). Stop-IRES contains the stop codon UGA at position 6 and has a peptide coding sequence designed to give a relatively high, detectable amount of readthrough even in the absence of NonSups. The design is based on studies showing that readthrough at the UGA stop codon proceeds in higher yields than at either the UAA and UAG stop codons^17^ and that such readthrough is further increased by both a downstream CUA codon (encoding Leu) at codon 7^18,19^ and an upstream AA sequence as part of the CAA codon 5 (encoding Gln).^17^ In Trp-IRES, UGA is replaced by UGG which is cognate to tRNA^Trp^, the most efficient natural tRNA suppressor of the UGA stop codon.^20,21^ Trp-IRES encodes the octapeptide FKVRQWLM, which permits facile quantification of octapeptide synthesis by ^35^S-Met incorporation. Although the mRNA attached to the CrPV-IRES contains additional codons downstream from the AUG encoding Met-tRNA^Met^, peptide synthesis is halted after Met incorporation by omitting the Thr-tRNA^Thr^ charged isoacceptor tRNA that is encoded by codon 9. Thus, the reaction terminates with FKVRQWLM-tRNA^Met^ bound to the ribosome P-site.

**Figure 2.**
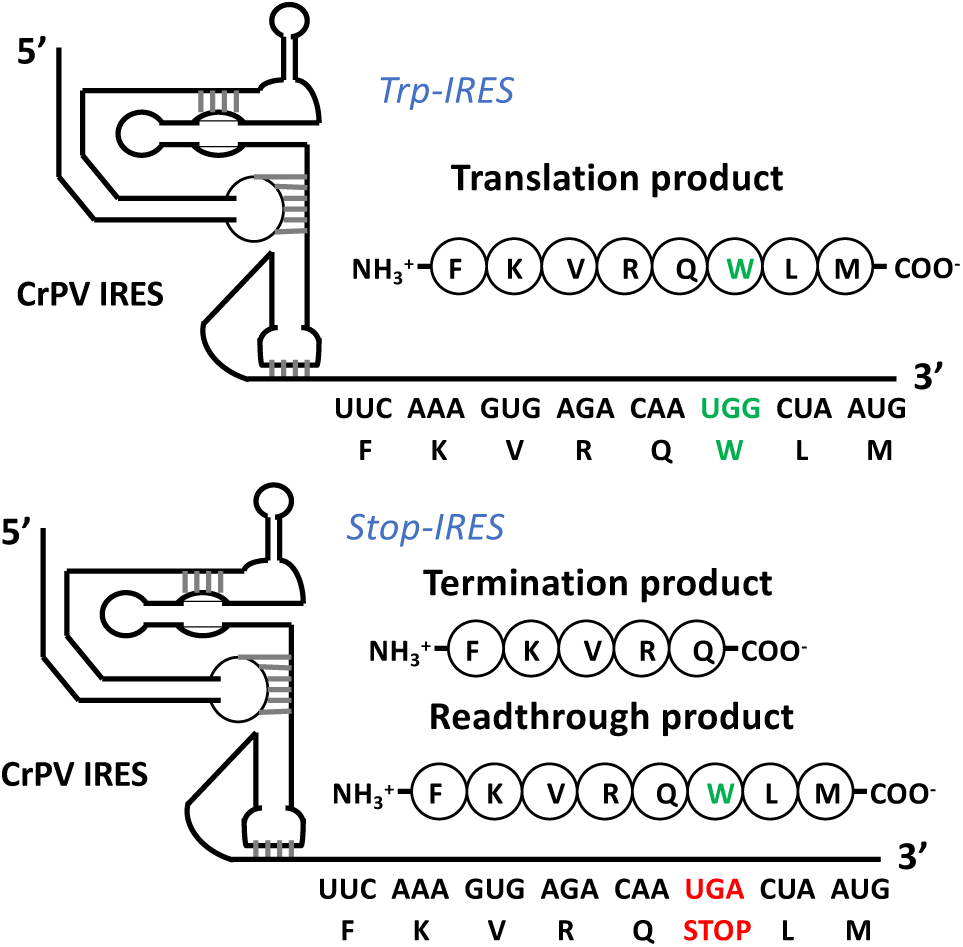
Coding sequences of Trp-IRES and Stop-IRES

For the results reported below, we prepared two POST5 translocation complexes, each containing FKVRQ-tRNA^Gln^ in the P-site, using ribosomes programmed with either Stop-IRES or Trp-IRES. The amounts of POST5 complex were quantified using [^3^H]-Gln. We next determined the amount of FKVRQWL[^35^S]-M-tRNA^Met^ that cosediments with the ribosome following incubation of each POST5 complex with a mixture of Trp-tRNA^Trp^, Leu-tRNA^Leu^, [^35^S]-Met-tRNA^Met^, elongation factors eEF1A and eEF2 and release factors eRF1 and eRF3. Octapeptide formation can also be quantified by hydrolyzing FKVRQWLM-tRNA^Met^ with strong base and measuring the released [^35^S]-labeled octapeptide following a thin layer electrophoresis purification.^14,22^ Although both methods gave very similar results (Figure S1), we prefer the cosedimentation assay because it is quicker and affords results having lower variability.

For Trp-IRES, synthesis of octapeptide from POST5 complex proceeded in high yield, giving a measured octapeptide/POST5 ratio of 0.75 ± 0.08 (n = 24). The amount of Stop-IRES readthrough is a sensitive function of release factor concentrations, decreasing dramatically as these concentrations are raised. We adjusted the release factor concentrations to give a basal octapeptide/POST5 ratio of 0.08 ± 0.02. This value permits facile estimation of NonSup concentrations we employ result in NonSup-induced readthrough efficiences which are high enough to make it feasible to perform experiments, currently underway, that will elucidate the detailed mechanisms of NonSup-induced readthrough.

Results measuring NonSup-induced readthrough of Stop-IRES are presented in Figure 3. The saturation curves obtained with the 7 aminoglycosides (AGs) examined (Figure 3A) are each consistent with a single tight site of AG binding to the ribosome which induces readthrough, with EC_50_s falling in the range of 0.4 – 4 μM and plateau octapeptide/POST5 ratios varying from 0.1 – 0.3 (Table S1) above the basal level measured in the absence of added NonSup (see *Supporting Information*, Item 2). These results are consistent with results on readthrough obtained in intact cells showing that a) G418, NB84, NB124^23^ and gentamicin X2^24^ are much more effective than the gentamicin mixture currently approved as an antibiotic; and b) NB84, NB124,^23^ and gentamicin X2^24^ have similar potencies, measured by either EC_50_ or readthrough efficiency. These consistencies suggest that aminoglycosides stimulate readthrough in cells primarily by direct effects on the protein synthesis machinery.

**Figure 3.**
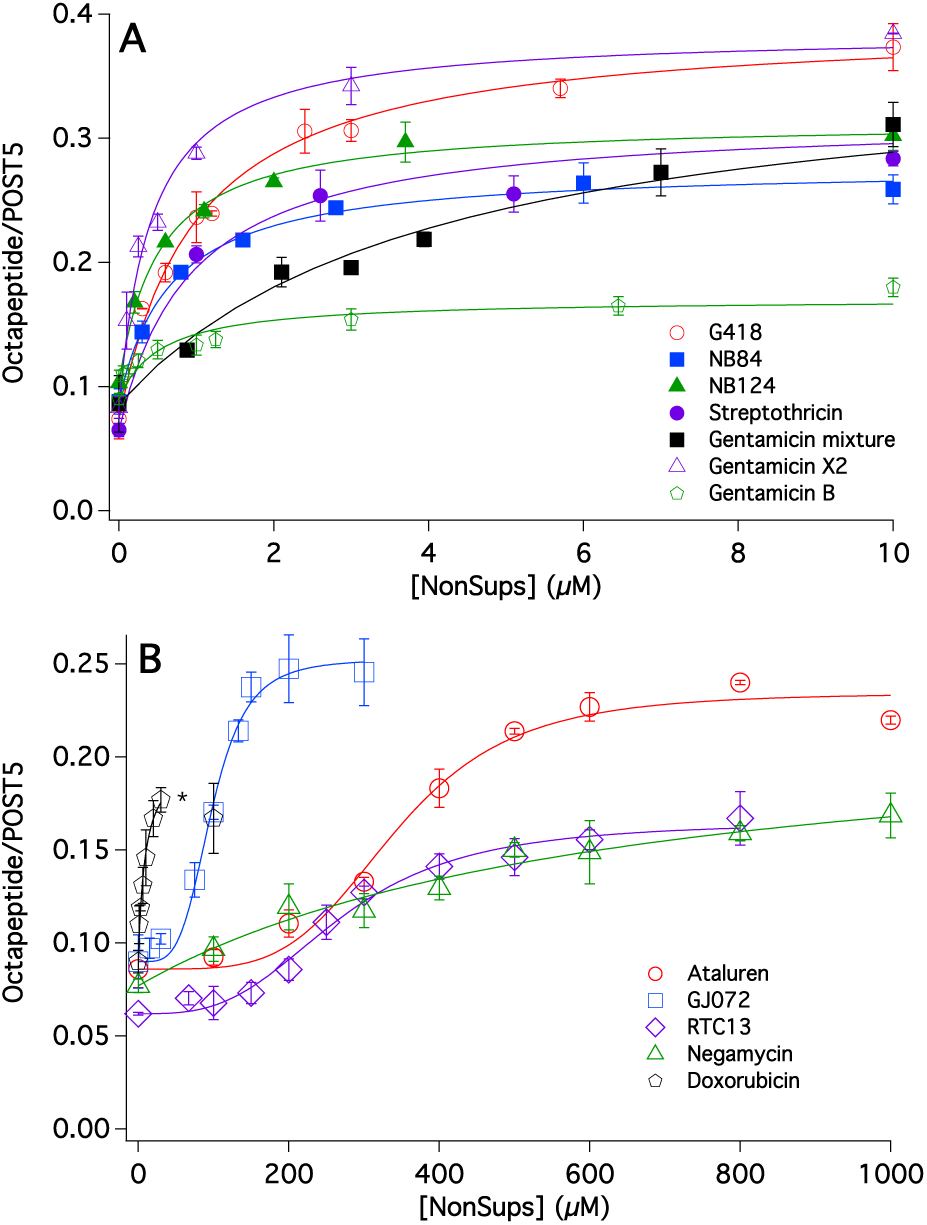
Readthrough as a function of nonsense suppressor concentration. **A.** Aminoglycosides. **B.** Ataluren-like NonSups and Others. None of the NonSups in Figure 3 showed appreciable inhibition of octapeptide formation from pentapeptide by ribosomes programmed with Trp-IRES at concentrations equal to twice their EC_50_ values. *The highest doxorubicin employed was 30 μM because higher concentrations led to significant ribosome and Met-tRNA^Met^ particle formation (Figure S7).

The NonSups ataluren, GJ072, and RTC 13 share similar structures, containing a central aromatic heterocycle having two or three substituents, at least one of which is aromatic induced enhancement of readthrough. At the same time, the low release factor (Figure 1). They also show similar S-shaped readthrough concentration-dependent activity curves (Figure 3B), with EC_50_ values between 0.17 – 0.35 mM and plateau octapeptide/POST5 ratios ranging from 0.10 – 0.16 (Table S1). These S-shaped curves yield Hill n values of ~ 4, which suggest multi-site binding of ataluren-like NonSups to the protein synthesis machinery. Formation of NonSup aggregates in solution which induce readthrough could also give rise to S-shaped curves, but we consider this to be unlikely based on the constancy of the chemical shift and line shape of ataluren’s ^19^F NMR peak over a concentration range of 0.03 – 2.0 mM (see *Supporting Information*, Item 6). Detailed dose-dependent readthrough results are available for live cell activities of ataluren^8^ and GJ072,^25^ each showing bell-shaped curves that contrast with the saturation behavior seen in Figure 3B. The decrease in readthrough activity at higher Non-Sup concentrations may reflect interactions of ataluren and GJ072 with cellular components not present in our *in vitro* assay.

Two other reported NonSups, negamycin^26^ and doxorubicin^27^, also display readthrough activity (Figure 3B). Both show a simple activity curve, with similar readthrough efficiencies (0.10 – 0.13) but a 50-fold difference in EC_50_ values, with doxorubicin having the lower value (Table S1). These compounds have potential for future development (see *Supporting Information*, Item 10).

Several other compounds that have readthrough activity in cellular assays, tylosin,^28^ azithromycin,^29^ GJ071^25^ and escin^27^ show little or no readthrough activity in our assay in the concentration range 30 – 600 μM (Figure S5). In addition, escin at high concentration inhibits both basal readthrough and normal elongation, the latter measured with Trp-IRES programmed ribosomes, with the effect on basal readthrough being much more pronounced (Figure S6). These results suggest that readthrough effects of tylosin, azithromycin, GJ071 and escin in cellular assays are likely to arise from interactions not directly involving the protein synthesis apparatus.

The assay results reported so far utilize a single time point, corresponding to full reaction leading either to octapeptide or pentapeptide formation (Figure 2). The clear difference between the saturation curves seen for aminoglycosides vs. ataluren-like NonSups raised the question of whether the kinetics of NonSup-induced octapeptide formation might also show differences. Accordingly, we compared the kinetics of octapeptide formation using ribosomes programmed with Trp-IRES and Stop-IRES in the presence of either G418 or ataluren. The results (Figure 4) show similar rate constants (Table S2) (1.0 – 1.6 min^−1^) for octapeptide formation by Trp-IRES in the presence or absence of G418 or ataluren, and by Stop-IRES in the presence of G418, but a much slower rate constant (0.14 min^−1^) for Stop-IRES in the presence of ataluren. Additional experiments, described in Supplementary Information, Item 3, strongly indicate that, as expected, this slow rate constant for ataluren-induced octapeptide formation is due to slow conversion of Stop-IRES POST5 complex to POST6 complex. These experiments also indicate that Stop-IRES POST6 complexes are labile, in contrast to the stable Trp-IRES POST6 complex (Figure S3). This lability is likely a consequence of imperfect base-pairing in the P-site between the anticodon loop of FKVRQW-tRNA^Trp^ and the UGA stop codon.

**Figure 4.**
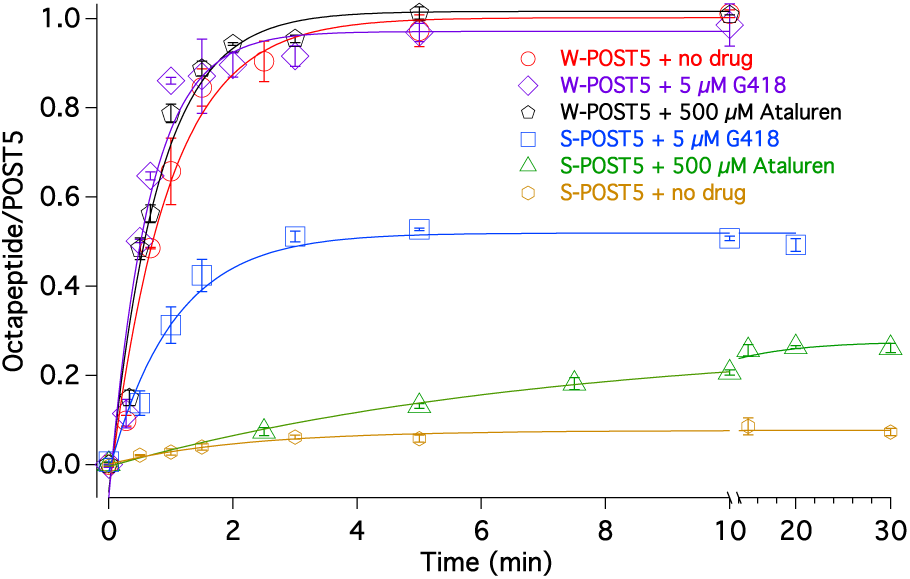
Kinetics of NonSup-induced octapeptide formation from Trp-IRES POST5 and Stop-IRES-POST5.

The results presented in Figures 3 and 4 suggest that aminoglycosides and ataluren-like compounds stimulate readthrough by different mechanisms, AGs via binding to a single tight site on the ribosome and ataluren-like compounds via weaker, multi-site binding which induces a slower change in the protein synthesis apparatus that permits readthrough.

EC_50_ values found in intact cells differ considerably from those measured by our cosedimentation assay, being much higher for AGs,^23,30^ and much lower for ataluren,^8,31^ RTC13^25^ and GJ072.^25^ We attribute these differences to the poor uptake of positively charged aminoglycosides into cells, while uptake is favored for the hydrophobic ataluren-like molecules. Thus, vis-à-vis the culture medium, intracellular concentration would be expected to be lower for AGs and higher for ataluren, RTC13 and GJ072.

Aminoglycosides have well-characterized tight binding sites in both prokaryotic^32^ and eukaryotic ribosomes,^33^ proximal to the small subunit decoding center, that have been linked to their promotion of misreading, although binding to additional sites at higher aminoglycoside concentrations have also been observed.^34^ Similarly, the functionally important prokaryotic ribosome binding site of negamycin has also been identified within a conserved small subunit rRNA region that is proximal to the decoding center,^32,35^ and it is not unlikely that this site is also present in eukaryotic ribosomes. However, nothing is known about the readthrough-inducing sites of action within the protein synthesis apparatus of the ataluren-like NonSups (Figure 3B) or of doxorubicin. Indeed, it has even been suggested that ataluren may not target the ribosome.^36^ Although aminoglycosides have been the subject of detailed mechanism studies of their effects on prokaryotic misreading (references provided in *Supporting Information*, Item 10) questions remain over their precise modes of action, and detailed mechanistic studies on aminoglycoside stimulation of readthrough and misreading by eukaryotic ribosomes are completely lacking. Virtually nothing is known about how negamycin, doxorubicin, and the ataluren-like NonSups stimulate eukaryotic readthrough.

Our laboratories are currently engaged in studies to elucidate the detailed mechanisms of action of NonSups directly interacting with the protein synthesis machinery. Such studies include smFRET measurements, which have provided detailed information about processive biochemical reaction mechanisms, particularly in the study of protein synthesis (references provided in *Supporting Information*, Item 10). Two fluorescent labeled tRNAs, when bound simultaneously to a ribosome, at either the A-and P-sites in a pretranslocation complex, or the P-and E-sites in a postranslocation complex, are spaced appropriately to generate a robust FRET signal.^37,38^ In preliminary work we have observed tRNA-tRNA FRET in the pretranslocation complex Trp-IRES-PRE6, which has tRNA^Gln^(Cy5) in the P-site and the peptidyl-tRNA, FKVRQW-tRNA^Trp^(Cy3) in the A-site (Figure 5A). The FRET efficiency of 0.47 ± 0.02 is similar to the value recently reported for neighboring tRNAs, also labeled at or near the elbow region of tRNA, bound in a PRE complex to the human ribosome.^34^ Addition of eEF2.GTP converts Trp-IRES-PRE6 to a Trp-IRES-POST6 complex, containing tRNA^Gln^(Cy5) in the E-site and FKVRQW-tRNA^Trp^(Cy3) in the P-site, which is accompanied by an increase in Cy3:Cy5 FRET efficiency to 0.73 ± 0.01 (Figure 5A). Repetition of this experiment with Stop-IRES-POST5 in the absence of eEF2 decreased the number of pretranslocation complexes (Stop-IRES-PRE6) formed to 22% of that seen with Trp-IRES. POST5 complexes that did not bind tRNA^Trp^-TC were readily distinguishable due to the lack of direct Cy3 and sensitized Cy5 emission (Figure 5B, top) but did display substantial emission and photobleaching when Cy5 was illuminated directly (Figure 5B, bottom). Baseline readthrough was increased by addition of G418 (1 – 10 μM) with a concentration dependence consistent with the EC_50_ value of 0.98 μM shown in Table S1. This agreement between the ensemble and single molecule assays demonstrates our ability to monitor NonSup-induced readthrough by smFRET, which, in subsequent studies, will allow determination of the effects of NonSups on the dynamics of the nascent peptide elongation cycle that commences with suppressor tRNA recognition of a premature stop codon.

**Figure 5.**
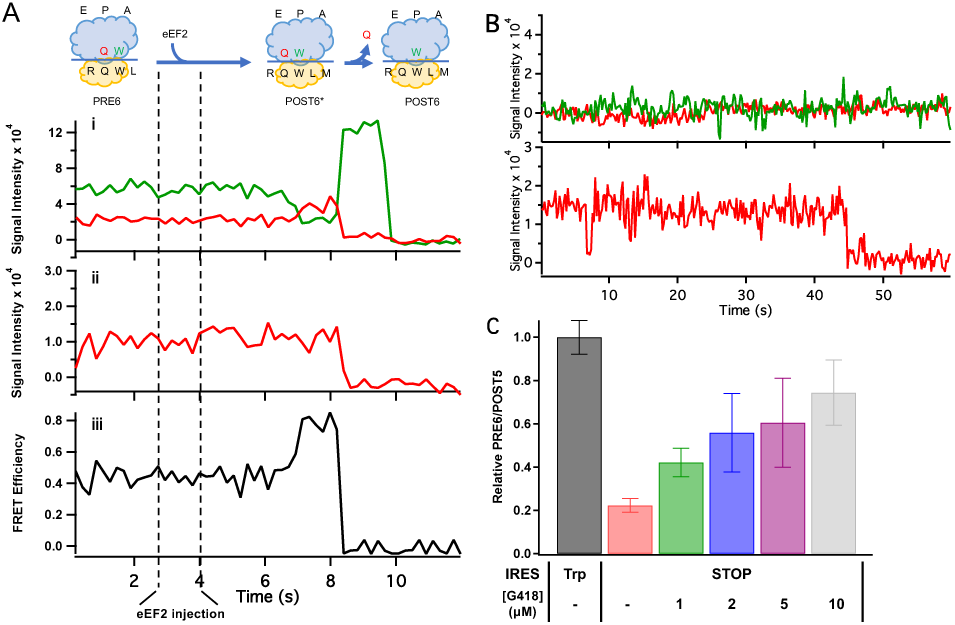
smFRET experiments. **A.** eEF2-induced translocation of the 80S-Trp-IRES-PRE6 complex to form 80S-Trp-IRES-POST6 complex followed by release of tRNA^Gln^. The cartoon at the top shows the state progression during translocation. i. Single molecule traces. Green and red traces show tRNA^Trp^(Cy3) emission and tRNA^Gln^(Cy5) sensitized emission, respectively, following eEF2 injection, excited at 532 nm. ii. ALEX intensity signal from direct excitation of tRNA^Gln^(Cy5) at 640 nm. iii. FRET efficiency between tRNA^Trp^(Cy3) and tRNA^Gln^(Cy5) showing a transient increase following eEF2 injection on conversion of the PRE6 complex to POST6. **B.** Recording of fluorescence traces for a Stop-IRES complex that did not bind tRNA^Trp^. Colors as in panel A. **C.** Effects of increasing G418 concentration on PRE6 complex formation from 80S-Stop-IRES-POST5 complex. Added G418 did not affect formation of Trp-IRES-PRE6 from Trp-IRES-POST5.

We expect the results of such smFRET studies, when combined with those of related ensemble mechanistic studies and studies directed toward identifying sites of NonSup interaction, to aid in achieving the understanding needed to improve the clinical efficacy of NonSups.

## Experimental procedures

### Materials

Shrimp (*A. salina*) ribosomes and ribosomal subunits, yeast elongation factors eEF2 and eEF1A, tRNAs and Trp-IRES and Stop-IRES, POST4 and POST5 complexes were prepared essentially as described in Zhang *et al*.^14^ Yeast 6xHis-tagged release factors (full-length eRF1 and amino acids 166-685 of eRF3), expressed in *E. coli* and purified using a TALON cobalt resin, were a generous gift from Alper Celik (University of Massachusetts Medical School). Nonsense suppressors (Figure 1) were obtained as follows: gentamicin mixture and G418 (Sigma), nourseothricin sulfate, a mixture of streptothricins D and F (Gold Biotechnology), doxorubicin (Fisher Scientific), escin and tylosin (Alfa Aesar), azithromycin (APExBIO), gentamicin B (MicroCombiChem). PTC Therapeutics supplied the following NonSups: ataluren sodium salt, RTC13, GJ071, GJ072, and gentamicin X2. Negamycin was a gift from Alexander Mankin, University of Illinois at Chicago. NB84 and NB124^23^, currently available as ELX-03 and ELX-02, respectively, from Eloxx Pharmaceuticals (Waltham, MA), were gifts from Timor Baasov (Technion, Haifa).

### Octapeptide formation readthrough assays

POST5 complex (0.02 μM) was mixed with Trp-tRNA^Trp^, Leu-tRNA^Leu^, and [^35^S]-Met-tRNA^Met^ (0.08 μM each), elongation factors eEF1A (0.08 μM), eEF-2 (1.0 μM) and release factors eRF1 (0.010 μM) and eRF3 (0.020 μM) and incubated at 37 °C in Buffer 4 (40 mM Tris-HCl pH 7.5, 80 mM NH_4_Cl, 5 mM Mg(OAc)_2_, 100 mM KOAc, 3 mM 2-mercaptoethanol) either for 20 min (Figure 3), or for times varying from 30 s to 20 min (Figure 4), in the absence or presence of NonSups. For octapeptide determination by co-sedimentation of FKVRQWL[^35^S]M-tRNA^Met^ with the ribosome, reaction mixture aliquots (40 μL) were quenched with 150 μL of 0.5 M MES buffer (pH 6.0) at 0 °C, which inhibits peptide bond formation. Following addition of carrier 70S *E. coli* ribosomes (100 pmol, 3 μL of 33 μM 70S), all ribosomes were pelleted by ultracentrifugation through a 1.1 M sucrose solution in Buffer 4 (350 μL) at 540,000 × g for 70 min at 4 °C. The ribosome pellet was resuspended in Buffer 4, and cosedimenting FKVRQWL[^35^S]M-tRNA^Met^ was determined. For octapeptide determination by thin layer electrophoretic purification of base-released FKVRQWL[^35^S]M, reaction mixture aliquots (80 μL) were quenched with 0.8 M KOH (9 μL of 8M KOH) and the base-quenched samples were incubated at 37 °C for 1 h. Acetic acid (9 μL) was then added to lower the pH to 2.8. Samples were next lyophilized, suspended in water, and centrifuged to remove particulates, which contained no ^35^S. The supernatant was analyzed by thin layer electrophoresis as previously described,^22^ using the same running buffer. The identity of FKVRQWLM was confirmed by the co-migration of the ^35^S radioactivity with authentic samples (Figure S1A) obtained from GenScript (Piscataway, NJ). The ^35^S radioactivity in the octapeptide band was used to determine the amount of octapeptide produced. Details concerning measurement of assay background and basal level of readthrough in the absence of added Non-Sup may be found in *Supporting Information*, Item 2.

### Other methods

smFRET experiments were performed essentially as described in Chen *et al*.^38^ Ataluren ^19^F NMR spectra were performed in buffer 4 with 10% D_2_O on a Bruker DMX 360 MHz NMR spectrometer with a 5 mm Quattro Nucleus Probe. Data were analyzed with mNova software.

## ASSOCIATED CONTENT

### Supporting Information

One pdf file containing additional experimental details and references (Items 1 −10), 7 Figures, 3 Tables. The *Supporting Information* is available free of charge on the ACS Publications website at DOI: XXX.

## Author Contributions

The manuscript was written through contributions of all authors. All authors have given approval to the final version of the manuscript.

## ACKNOWLEDGMENTS

We thank Drs. Alexander Mankin (University of Illinois at Chicago, negamycin), Timor Baasov (Technion, Haifa, NB84, NB124) and Alper Celik (University of Massachusetts Medical School, eRF1, eRF3) for reagents used in these studies and Carla Zimmerman for technical assistance. This work was supported by research grants to BSC (PTC Therapeutics; the Orphan Disease Center, University of Pennsylvania MDBR-17-108-CF1282X; Cystic Fibrosis Foundation-COOPER18G0; NIH GM127374) to YEG (NIH GM118139), to MR (Michael Smith Foundation for Health Research) and AJ (NIH GM27757 and GM122468).

## ABBREVIATIONS

AG: aminoglycoside
CrPV: cricket paralysis virus
IRES: internal ribosome entry site
NonSups: nonsense suppressors
PTC: premature termination codon
smFRET: single molecule fluorescence resonance energy transfer

